# SMAC: identifying DNA N^6^-methyladenine (6mA) at the single-molecule level using SMRT CCS data

**DOI:** 10.1101/2024.11.13.623492

**Authors:** Haicheng Li, Junhua Niu, Yalan Sheng, Yifan Liu, Shan Gao

**Affiliations:** MOE Key Laboratory of Evolution & Marine Biodiversity and Institute of Evolution & Marine Biodiversity, Ocean University of China, Qingdao 266003, China; Laboratory for Marine Biology and Biotechnology, Qingdao Marine Science and Technology Center, Qingdao 266237, China; Shum Yiu Foon Shum Bik Chuen Memorial Centre for Cancer and Inflammation Research, School of Chinese Medicine, Hong Kong Baptist University, Hong Kong, SAR, China; Department of Biochemistry & Molecular Medicine, University of Southern California Keck School of Medicine, Los Angeles, CA 90033, USA

**Keywords:** DNA N^6^-methyladenine (6mA), single molecule, SMRT CCS sequencing

## Abstract

DNA modifications, such as N^6^-methyladenine (6mA), play important roles in various processes in eukaryotes. Single molecule, real-time (SMRT) sequencing enables the direct detection of DNA modifications without requiring special sample preparation. However, most SMRT-based studies of 6mA rely on ensemble-level consensus by combining multiple reads covering the same genomic position, which misses the single-molecule heterogeneity. While recent methods have aimed at single-molecule level detection of 6mA, limitations in sequencing platforms, resolution, accuracy, and usability restrict their application in comprehensive epigenetic studies. Here, we present SMAC (Single Molecule 6mA analysis of CCS reads), a novel framework for accurately detecting 6mA at the single-molecule level using SMRT CCS data from the Sequel II system. It is an automated method that streamlines the entire workflow by packaging both existing software and built-in script, with support for user-defined parameters to allow easy adaptation for various studies. This algorithm utilizes the statistical distribution characteristics of enzyme kinetic indicators to identify 6mA of each DNA molecule, rather than relying on a fixed cutoff, which significantly improves accuracy at the single-nucleotide and single-molecule level. SMAC is a powerful new tool that enables de novo detection of 6mA and empowers investigation of its functions in modulating physiological processes.

## Introduction

DNA modifications, such as N^6^-methyladenine (6mA), play significant roles in various crucial processes in eukaryotes, including DNA structure maintenance, transcriptional regulation, nucleosome positioning, transposon activation, and DNA replication. It plays a vital biological role in stress responses, embryonic development, cellular physiological states, tumor cell growth, plant growth and development, as well as immune responses (1–12). Understanding the distribution of 6mA in eukaryotic genomes is essential for uncovering its regulation mechanisms and biological implications.

Several methodologies have been developed to detect 6mA, including dot blot, liquid chromatography-mass spectrometry (LC-MS), and 6mA immunoprecipitation sequencing (6mA-IP-seq) (13–15). While these techniques have provided valuable insights, each comes with specific limitations. For instance, dot blot and LC-MS can estimate total 6mA levels, but they provide no information of context sequences and cannot rule out contamination from DNA of other species (13,15). Additionally, 6mA-IP-seq, which relies on Illumina short-fragment sequencing, can identify 6mA-enriched genomic regions but lacks single-nucleotide resolution (13–15). Moreover, both dot blot and 6mA-IP-seq may produce false-positive signals due to non-specific antibody binding(13,15). In contrast, Single molecule, real-time (SMRT) sequencing offers the direct detection of DNA modifications at the single-base resolution across long genomic regions (13–15), making it a powerful tool for detecting 6mA with unprecedented accuracy and detail.

During SMRT sequencing, double-stranded native DNA fragments are circularized by ligating hairpin adapters to each end. DNA polymerase then proceeds around the circularized DNA multiple times, with the number of passes depending on the fragment size and polymerase processivity (16,17). The time taken by the polymerase to translocate from one nucleotide to the next is termed the inter-pulse duration (IPD), and variations in IPDs are highly correlated with modifications in the DNA template (17–19). SMRT sequencing can be performed in two modes: Continuous Long Read (CLR) and circular consensus sequencing (CCS). While CLR is effective for mapping 6mA at the ensemble level by combining data from different DNA molecules covering the same genomic position, it lacks the ability to provide single-molecule information. In contrast, CCS, with shorter fragment sizes and improved polymerase processivity, enables more passes (subreads) over the same DNA molecule, thus generating high-fidelity (HiFi) reads with improved sequence accuracy (20–22). More importantly, subreads are combined to accurately call 6mA at single-molecule levels, allowing for exploration of heterogeneities among different molecules (23–25).

The classic standard SMRT-seq pipeline, such as the ipdSummary tool in SMRT Link software, can be applied to both CLR and CCS but has typically been used to detect 6mA at the ensemble level (17,26). While frameworks like SMALR have been developed for single-molecule 6mA detection, they were designed for earlier CCS data produced by the PacBio RSII system (23). In addition, its case-control method requires sequencing methylation-free samples, significantly increasing costs. With advancements in PacBio SMRT CCS technology, the datasets generated by the Sequel II system have increased tenfold in size. More recent CCS-based methods, such as 6mASCOPE, offer higher sensitivity but do not output results at the single-nucleotide resolution, instead providing only an assessment of the overall 6mA/A level (27). Another approach, 6mA-Sniper, enables single-molecule 6mA detection but relies on fragmented scripts and has not been tested on species other than *Caenorhabditis elegans* (28). Due to limitations in current 6mA detection tools and pipelines base on CCS data, many studies still rely on the CLR data, or CCS data but analyzed using standard SMRT-seq pipeline (7,11,12,29–33), failing to capture the 6mA methylation states of individual molecules.

In this study, we present SMAC (<u>S</u>ingle <u>M</u>olecule 6m<u>A</u> analysis of <u>C</u>CS reads), a novel and automated pipeline designed for single-molecule, single-nucleotide, and strand-specific detection of 6mA using SMRT CCS data. SMAC is built on the continuously updated PacBio SMRT Link software and is compatible with the commonly used SMRT sequencing platform. Unlike SMALR that requires additional methylation-free datasets, SMAC employs *in silico* controls embedded in ipdSummary. It uses molecule-specific IPD ratio information to infer methylation states at the single-molecule resolution. SMAC offers several key advantages: First, it applies rigorous data pre-treatment to minimize background noise. Second, it identifies more reliable genome-wide 6mA sites than the standard SMRT-seq pipeline while also detecting 6mA at the single-molecule level. Third, it enhances the accuracy of 6mA detection within ApT motifs, where methylation can occur fully or hemi-methylated. Lastly, by using a Gaussian distribution fitting approach, SMAC allows for a more objective determination of the cutoff for 6mA site detection. SMAC is freely available on GitHub (https://github.com/liiihc/SMAC).

## Methods

SMAC includes multiple steps such as data preprocessing, alignment, and 6mA identification. The raw subreads data are processed using the ‘ccs’ module in SMRT Link to generate HiFi reads with the parameter ‘--hifi-kinetics’ and converted to FASTA format using the ‘bam2fasta’ module, which also provides a quality report for the sequencing run. By default, only reads with ≥ 20× passes are retained for downstream analysis. Users can modify the minimal number of passes by adjusting the “-passes” parameter within the SMAC pipeline, depending on their need for either larger datasets or stricter quality control. The HiFi reads are split into individual FASTA files to serve as reference sequences. The raw subreads are converted to SAM format using SAMtools and further split into individual SAM files for analysis. A reference index of the split HiFi reads is built using the ‘pbindex’ module in SMRT Link. Each SAM file is converted to BAM format and aligned to the corresponding HiFi reads using the ‘pbmm2’ module with the default parameter, while the IPD (inter-pulse duration) ratio is calculated using the ‘ipdSummary’ module.

The HiFi reads are aligned to the reference genome using both BLASTN and ‘pbmm2’, with only reads meeting the criteria of ≥ 80% coverage and ≥ 80% identity in the BLASTN results being retained for further analysis by default. Users can customize the BLASTN coverage and identity thresholds by modifying the “-coverage” and “-identity” parameter in the SMAC pipeline. To ensure accuracy, the IPD ratios of bases within 25 bp of the adapter sequences are trimmed by default. Users can modify the number of trimmed bases based on the IPD ratio variation in their own dataset, by adjusting the “-trimmer” parameter in the SMAC pipeline. Each adenine (A) in the CCS reads is mapped back to the genome based on the ‘pbmm2’ alignment results. The IPD ratio distribution of all adenines aligned to the reference genome is calculated, and a Gaussian distribution is fitted to determine the initial cutoff. By default, only reads with standard deviation of IPD ratios ≤ 0.6 for non-6mA bases on both Watson and Crick strands are retained for downstream analysis. Users can modify the cutoff of standard deviation of IPD ratios based on the IPD ratio variation in their own dataset, by adjusting the “-sd_cutoff” parameter in the SMAC pipeline. For ApT dinucleotides, a second Gaussian fitting is applied to the IPD ratios of the corresponding adenine within the initially identified 6mApT sites to determine a secondary cutoff. Using these cutoffs, 6mA sites are distinguished from non-methylated adenines at the single-molecule level. Users can modify the “-ipd_cutoff” parameter to apply cutoff determined by the Gaussian fit or any specific cutoff. The penetrance of each 6mA site is defined as the ratio of the number of 6mA sites to the total number of adenines across all reads.

Users are required to have BLASTN, SMRT Link, perl module “Statistics::Descriptive” and python packages including “logomaker lmfit, scipy, statsmodels, pandas, matplotlib and numpy” installed in advance.

## Results and Discussion

### Conceptual view of SMAC

Figure 1 outlines the conceptual framework of SMAC, which uses the IPD ratio from the CCS mode of the Sequel II system to call 6mA at the single-molecule level. The process starts with generating high-fidelity (HiFi) reads from raw subreads data using the ccs module in SMRT Link (Pacific Biosciences). Subreads and HiFi reads are then split based on single-molecule IDs into single molecules, filtering out those with low passes. Next, subreads are aligned to their corresponding HiFi reads using the pbmm2 module in SMRT Link. The IPD ratios for all adenine sites are computed using the ipdSummary module in SMRT Link. Before calling 6mA, HiFi reads are mapped to the reference genome using BLASTN and pbmm2 to filter out reads originating from contaminants and locate the bases within the genome. A Gaussian distribution is then fitted to separate the 6mA peak from the background, establishing a cutoff value of IPD ratio to identify 6mA sites and assess the methylation states of 6mA at A on a single-molecule basis. During the data processing, SMAC provide quality control reports for base trimming, IPD ratio stability and basic statistics for library size and polymerase processivity. The final output includes 6mA calling results at both ensemble and single-molecule levels, along with the methylation states for each individual ApT dinucleotide. Most analyses presented in this study were based on native genomic DNA samples from the wild-type (WT, SB210 strain) *Tetrahymena thermophila*, an important unicellular model eukaryote with abundant 6mA in its genome (4,6,7,10,34,35), unless otherwise noted.

**Fig. 1.**
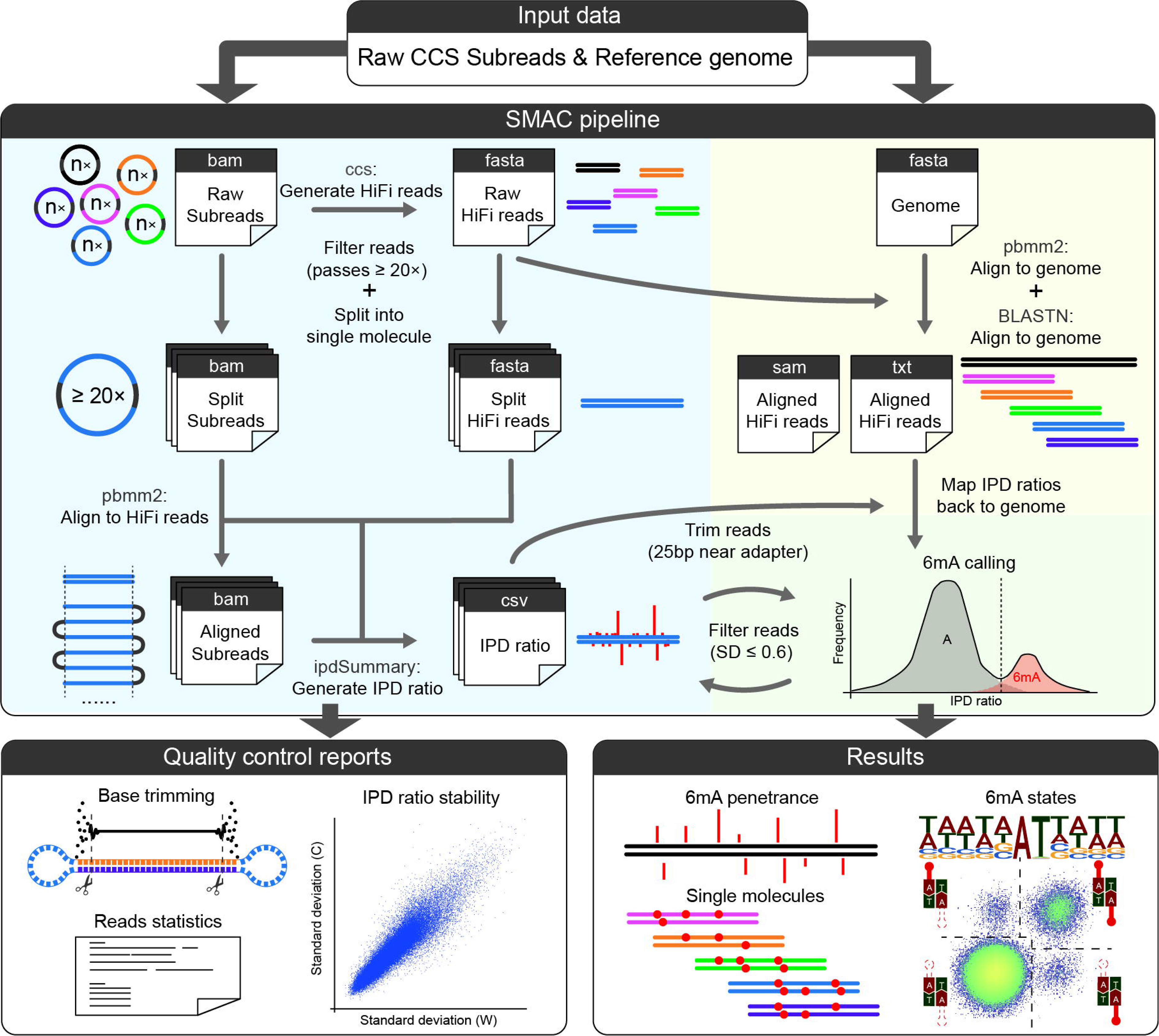
Conceptual view of SMAC (<u>S</u>ingle <u>M</u>olecule 6m<u>A</u> analysis of <u>C</u>CS reads). Overview of SMAC for detecting 6mA from SMRT CCS data. SMAC is an automated pipeline that processes original subreads files and the reference genome file of the target species, generating quality control reports and results of both ensemble and single-molecule level 6mA sites, along with the methylation states for each individual ApT dinucleotide. CCS, circular consensus sequencing. IPD, inter-pulse duration. Penetrance, the proportion of molecules containing 6mA at a specific genomic position.

SMAC has packaged all required steps into a comprehensive and flexible pipeline, enabling users to obtain analysis results simply by providing the original subreads files and the reference genome file of the target species. SMAC also allows users to adjust key parameters according to their own needs.

### Data trimming and filtration

SMRT CCS sequencing involves circular sequencing of SMRTbell templates, which consist of DNA molecules with adapters attached at both ends. To investigate whether the presence of these adapters can influence the IPD ratios near the ends of DNA molecules, we analyzed a published CCS dataset of whole genome amplification (WGA) for WT *T. thermophila* (10). By examining the IPD ratio distribution within a defined distance from the molecule ends, we observed increased variability in the bases within 25 bp of adapters (Figure 2A). Additionally, abnormal bases with an IPD ratio of zero were frequently observed in this region for unknown reasons (Figure S1A). Based on these findings, SMAC recommends trimming 25 bp from both ends of DNA molecules to minimize interference as a default setting.

**Fig. 2.**
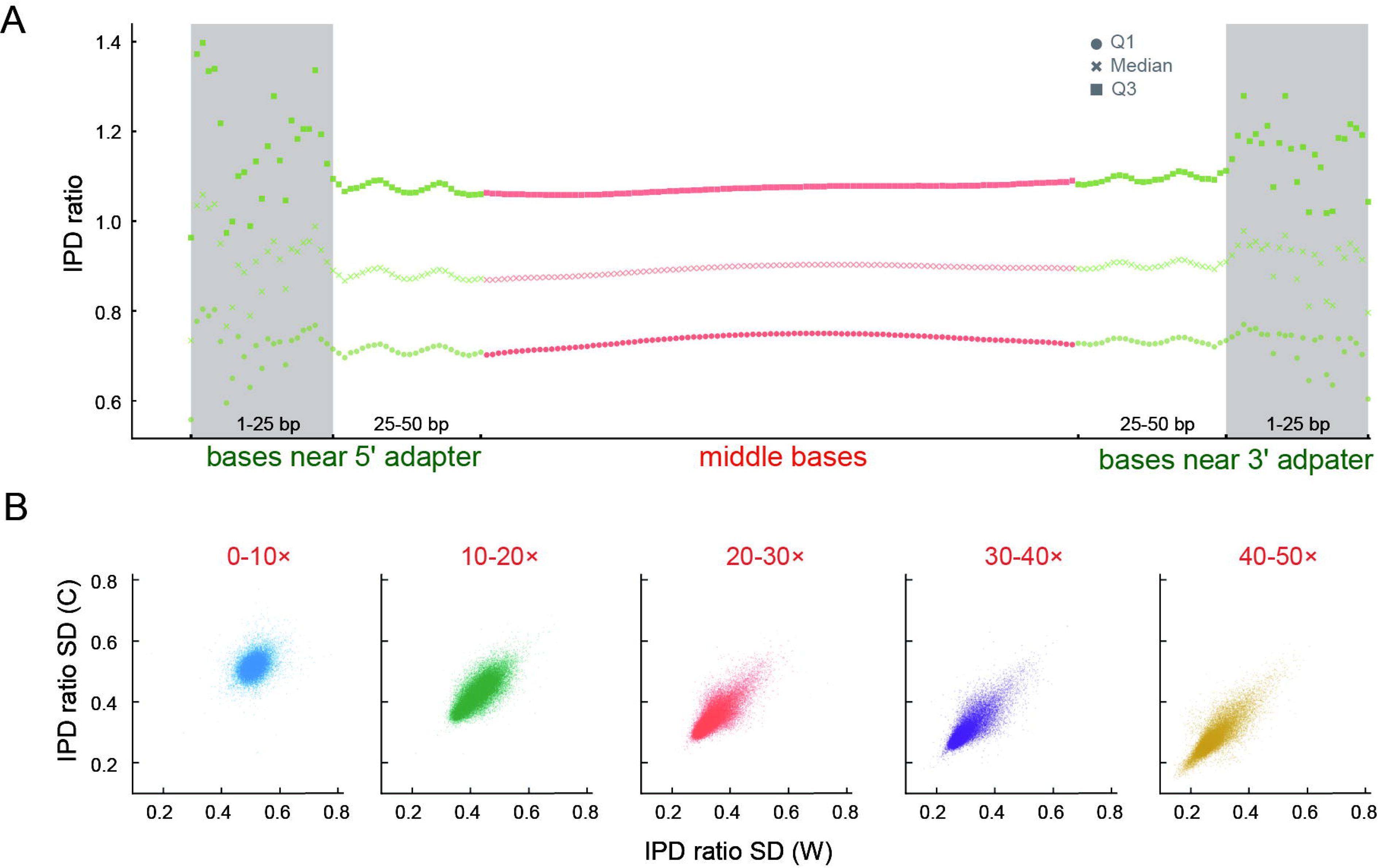
Data trimming and filtration. **A.** Distribution of IPD ratios in SMRT CCS reads. IPD ratios on bases within 50 bp near the 5’ or 3’ adapter were shown as green dots. Other bases located in the central region were normalized to a scale of 100 units and represented by yellow or orange? dots. Q1, median, and Q3 represent the first quartile, median, and the third quartile of IPD ratios. **B.** Average standard deviation (SD) of IPD ratios for SMRT CCS reads across different pass ranges: 0-10×, 10-20×, 20-30×, 30-40×, and 40-50×. SD values were calculated separately for IPD ratios on the Watson strand (W) and Crick strand (C).

The number of passes, or the number of subreads for each DNA molecule, significantly impacts the stability of IPD detection, with higher pass numbers resulting in more reliable IPD ratios. We categorized all molecules based on their number of passes and calculated the standard deviation of IPD ratios for bases on both the Watson and Crick strands, for the WGA sample. As the number of passes increased, the standard deviation decreased, stabilizing at about 20 passes (Figure 2B). Therefore, SMAC recommends filtering out molecules with fewer than 20 passes as a default setting.

Contamination from external DNA poses a significant challenge for 6mA detection, including mass spectrometry and dot blot assays. Even in SMRT sequencing, a few chimeric DNA molecules formed during library preparation can introduce considerable contamination, especially from bacterial DNA with high levels of 6mA. To address this, SMAC employs BLASTN to filter out chimeric molecules. We used a mixed sample of *Saccharomyces cerevisiae* (target species), *Escherichia coli* (contamination), and *Helicobacter pylori* (contamination) to examine the effect of BLASTN on reads mapping ratio and chimeric reads ratio. As the requirements for coverage and identity in BLASTN increased, the chimeric reads ratio gradually decreased, but the ratio of mapped reads retained after BLASTN alignment also significantly declined (Figure S1B). To minimize the proportion of chimeric reads while retaining as much valid data as possible, we established default BLASTN filtering parameters of 80 for both coverage and identity.

### Sensitive and accurate 6mA calling

In SMRT sequencing, 6mA exhibits higher IPD ratios compared to unmethylated adenines (A), leading to a bimodal distribution of IPD ratios in native DNA sample. In contrast, WGA sample lacking methylation display a single peak corresponding to A (Figure 3A). SMAC fits a Gaussian distribution to the high IPD ratio peak for native DNA samples, with the intersection of this fitted curve and the residual curve serving as the cutoff for 6mA detection (Figure 3A). Sites with IPD ratios exceeding this cutoff are identified as 6mA.

**Fig. 3.**
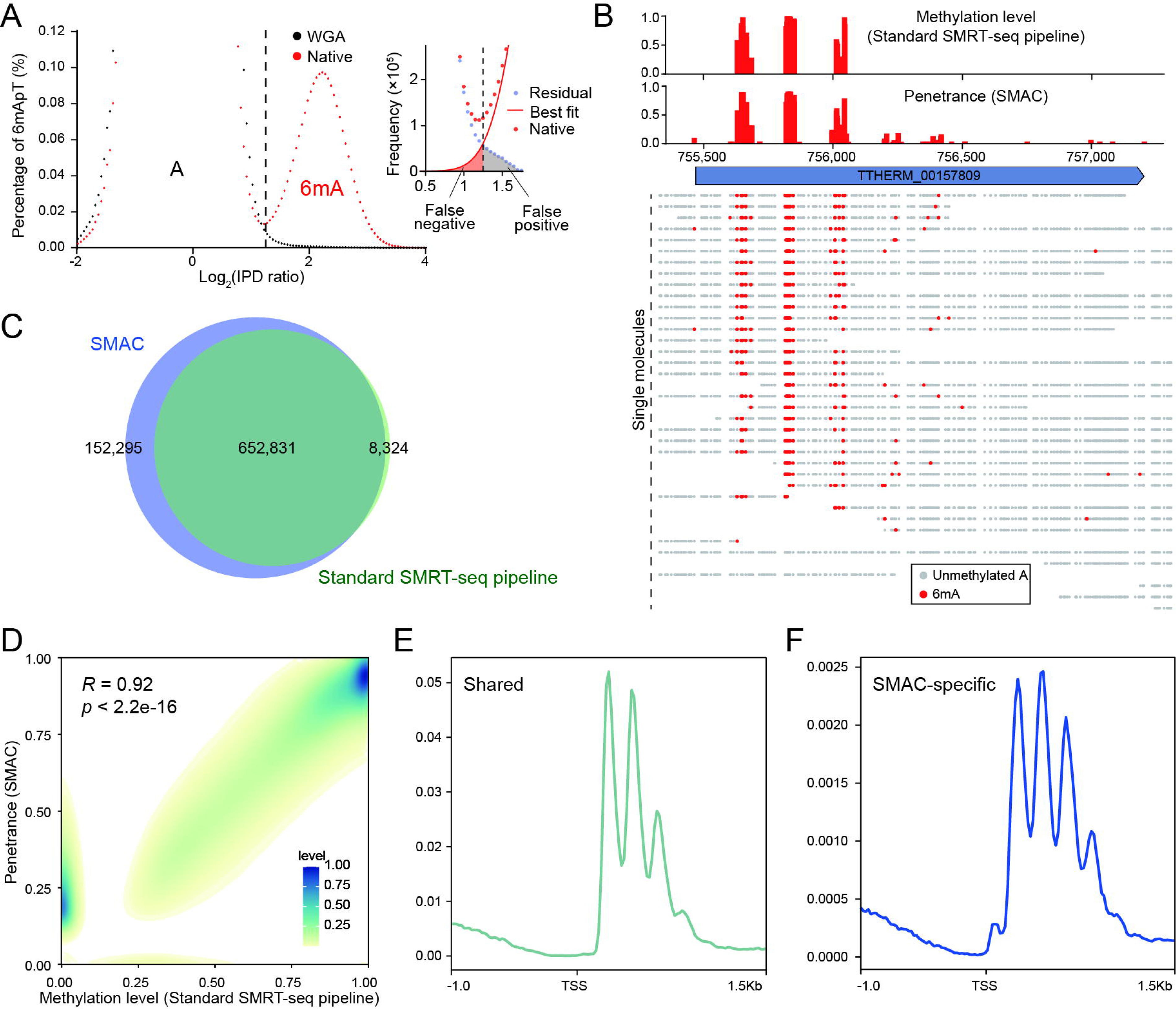
Sensitive and accurate 6mA calling. **A.** IPD ratio distribution (log_2_) of A sites in native DNA (red dots) and WGA (black dots) samples. Note the deconvolution of the 6mA peak (right) and the unmodified A peak (left) in IPD ratio distribution. Residual (light blue dots) represented the difference between authentic IPD distribution and the best fit curve of Gaussian distribution in the small 6mA peak (red line). The IPD ratio cutoff was indicated by dashed line. **B.** Comparison of penetrance the 6mA methylation level calculated by the standard SMRT-seq pipeline and penetrance calculated by SMAC, showcased by a representative genomic region from the native DNA sample. 6mA (red dots) and unmodified A sites (gray dots) on each individual DNA molecules were also displayed. **C.** Overlap between 6mA sites called by SMAC (6mA coverage ≥ 5x) and standard SMRT-seq pipeline (coverage ≥ 100, Qv ≥ 30). **D.** Correlation between 6mA methylation level calculated by standard SMRT-seq pipeline and penetrance calculated by SMAC. 6mA site distribution density was plotted as a heat map. **E.** Distribution profiles of 6mA sites identified by both standard SMRT-seq pipeline and SMAC. 6mA peaks had a periodicity distribution downstream of TSS. TSS, transcription start site. **F.** Distribution profiles of SMAC specific 6mA sites.

SMAC aligns sequences of individual molecules back to the genome and calculates the penetrance, which represents the proportion of molecules containing 6mA at a specific genomic position (Figure 3B). In native DNA samples, SMAC detected 652,831 shared 6mA sites with the standard SMRT-seq pipeline based on SMRT Link, accounting for 81.02% of total 6mA sites identified by SMAC (Figure 3C). Additionally, SMAC identified 152,295 unique sites (18.98%) that the standard SMRT-seq pipeline failed to detect (Figure 3C). Conversely, the standard SMRT-seq pipeline detected only 8,324 unique sites (1.03% of total 6mA sites identified by SMAC and standard SMRT-seq pipeline) (Figure 3C), demonstrating an overall higher 6mA/A level calculated by SMAC (SMAC: 0.74%, standard SMRT-seq pipeline: 0.63%). For the WGA sample lacking a bimodal distribution, SMAC identified no valid 6mA sites, as there was no 6mA peak for Gaussian fitting. In contrast, the standard SMRT-seq pipeline falsely identified 9,028 sites as 6mA (Figure S2A). Together, SMAC identified more high-confidence 6mA sites while maintaining a lower false positive rate.

Regarding the assessment of 6mA levels, the penetrance calculated by SMAC correlated well with the methylation level determined by the standard SMRT-seq pipeline for shared 6mA sites. Although SMAC-specific sites displayed relatively lower penetrance (Figure 3D, Figure S2B), they demonstrated high coverage, reflecting reliable sequencing depth (Figure S2B). More importantly, these SMAC-specific sites exhibited typical features of authentic 6mA, as demonstrated by shared 6mA sites. First, they had a strong preference for the ApT dinucleotide (Table S([0-9]+)). Second, they were predominantly located downstream of transcription start sites (TSS) and displayed a periodic distribution between nucleosome arrays (Figures 3E, 3F, Figure S2D) (4,6,7). As a control, those false positive sites in the WGA sample identified by the standard SMRT-seq pipeline also had high coverage but lacked these features (Figure S2C, S2D).

Based on these comparisons, we conclude that SMAC can identify authentic 6mA with higher sensitivity and accuracy than the standard SMRT-seq pipeline.

### Determination of 6mApT states

In most prokaryotes and unicellular eukaryotes, 6mA typically occurs at the self-complementary ApT dinucleotide, existing in either a fully methylated or hemi-methylated state (1,4,12,18,27,36,37). These states may have distinct downstream regulatory functions, highlighting the importance of accurately distinguishing between them (11,12,32). SMAC performs an initial Gaussian fitting of IPD ratios for all A bases to generate the first cutoff for calling methylated ApT (6mApT) (Figure 4A). Moreover, since the IPD ratio distribution pattern of A on the opposite strand of 6mApT identified after the initial Gaussian fitting slightly changed, the IPD ratio distribution of the opposing A within initially identified 6mApT sites is recalculated. A second Gaussian fitting is then conducted to generate the second cutoff to refine the 6mApT calling (Figure 4A). After two rounds of 6mA calling, ApT are classified into four groups: fully methylated, hemi-methylated on the Watson strand (hemi-W), hemi-methylated on the Crick strand (hemi-C), and unmethylated (Figure 4B).

**Fig. 4.**
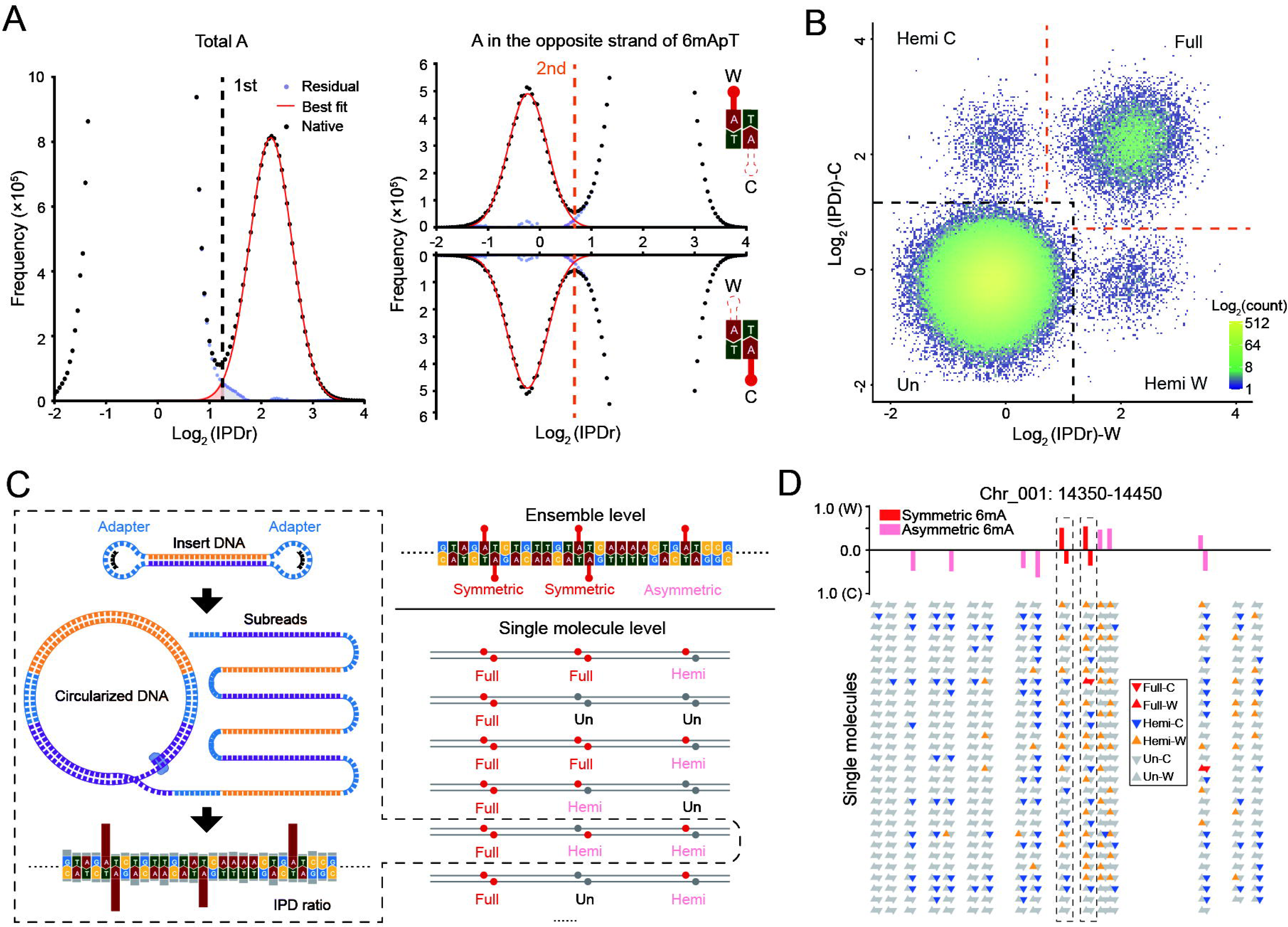
Determination of 6mApT states. **A.** Deconvolution of the 6mA peak and the unmodified A peak for IPD ratio distribution. The left panel showed the initial Gaussian fitting of the small 6mA peak of all A. The right panel showed the second Gaussian fitting result of the small unmodified A of the opposing A within initially identified 6mApT. Residual (light blue dots) was the difference between authentic IPD distribution and the best fit curve (red dots). The IPD ratio cutoff was marked by dashed line. **B.** Demarcation of the four methylation states of ApT duplexes by their IPD ratio on Watson (W) and Crick (C) strands, respectively. 6mA site distribution density was plotted as a heat map. The IPD ratio cutoff for 6mA calling was set according to deconvolution based on two rounds of Gaussian fitting. The IPD ratio cutoff was marked by dashed line. **C.** Schematic diagram illustrating the determination of 6mApT states at ensemble and single-molecule levels. The left panel showed the 6mA detection of a CCS single DNA molecule. The right panel showed the discrepancy between full/hemi sites at the single molecule level and symmetric/asymmetric methylation at the ensemble level. **D.** Representative DNA molecules from the maintenance methyltransferase knockout Tetrahymena strain Δ*AMT1*. The 6mA states as symmetric or asymmetric at the ensemble level identified by the standard SMRT-seq pipeline (top) were largely different from the full or hemi states revealed by SMAC (bottom). ApT dinucleotides with distinct methylation states (colored triangles) interspersed with unmodified ApTs (gray triangles).

Full and hemi-methylation states at the single-molecule level correspond to symmetric and asymmetric methylation at the ensemble level, respectively. However, the standard SMRT-seq pipeline tends to overestimate symmetric methylation by misinterpreting multiple hemi-methylated molecules as symmetric, whereas SMAC, by leveraging single-molecule information, avoids this issue (Figure 4C). We used both SMAC and the standard SMRT-seq pipeline to compare two *T. thermophila* samples: one predominantly featuring full methylation (WT, 89.29% full-6mApT) and another primarily exhibiting hemi-methylation (maintenance methyltransferase knockout strain Δ*AMT1*, 97.63% hemi-6mApT) (Table S([0-9]+)) (10). In both samples, the standard SMRT-seq pipeline revealed a much higher proportion of symmetric methylation compared to the full methylation level identified by SMAC, a discrepancy particularly pronounced in the sample dominated by hemi-methylation (Figure 4D, Table S([0-9]+)). Overall, SMAC outperforms the standard SMRT-seq pipeline in accurately determining 6mApT states.

### Detection limit and minimal sequencing depth

SMAC determines the threshold for 6mA detection by fitting a Gaussian curve to the bimodal distribution of IPD ratios. This strategy may be too stringent for samples with low methylation levels, as they might not display a clear bimodal pattern. To explore this issue, we diluted a native DNA sample (6mA/A= 0.74%) with a WGA sample to various levels and applied the bimodal model for curve fitting. The error rate for calling 6mA increased significantly when the 6mA/A level approached approximately 0.1%. When the 6mA/A level dropped below 0.04%, the IPD ratios of A no longer exhibited a bimodal distribution, with the 6mA signal on the right side completely masked by the noise from A on the left side (Figure 5A). Therefore, SMAC can reliably detect 6mA in samples where 6mA/A levels exceed 0.04%.

**Fig. 5.**
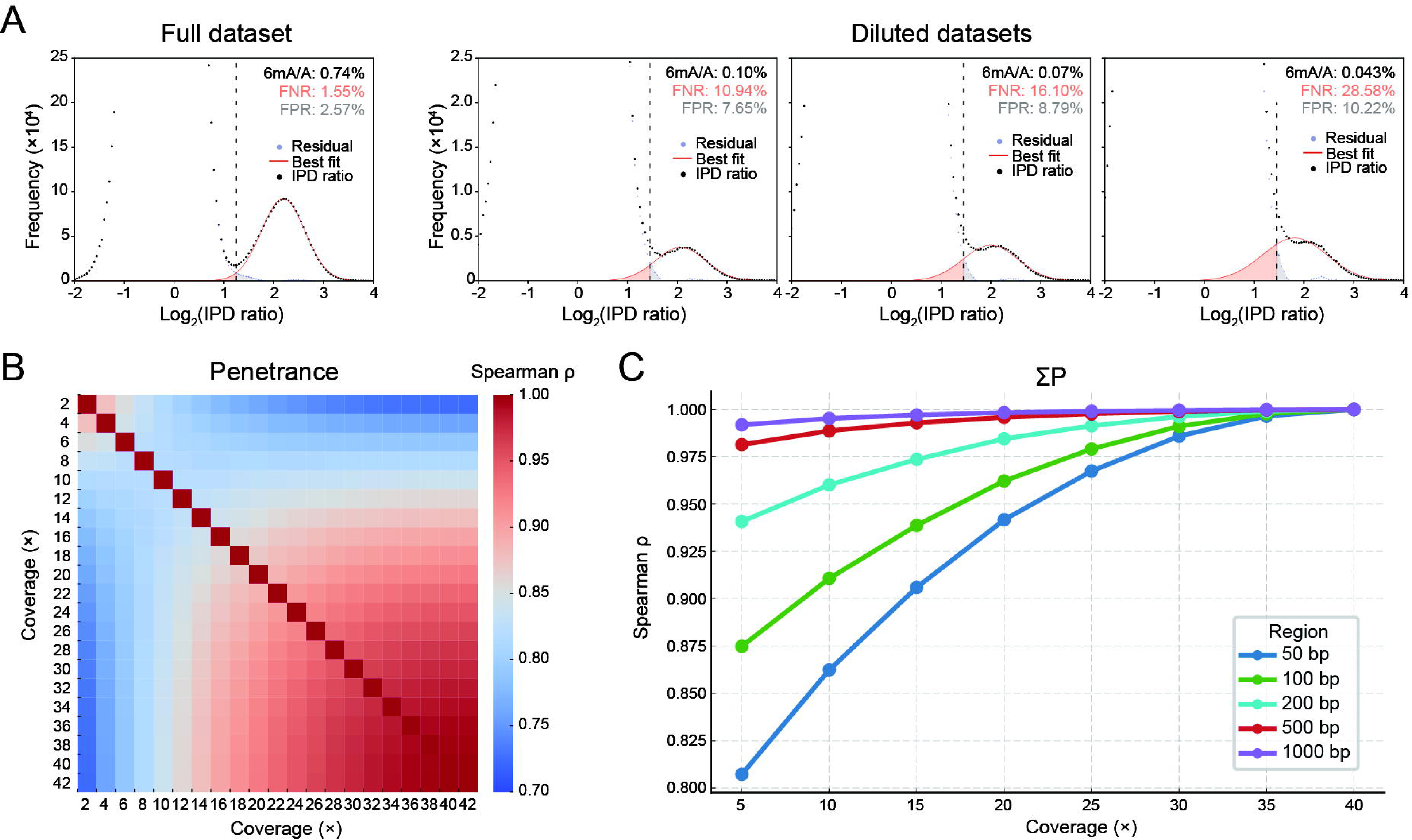
Impact of 6mA levels and sequencing coverage on reliable 6mA detection. **A.** Deconvolution of the 6mA peak and unmodified A peak for IPD ratio distribution in diluted datasets of native DNA and WGA. The native DNA data was mixed with WGA data with the ratios of data size at 1:6, 1:9, 1:15. Residual is the difference between authentic IPD distribution and the best fit curve (light blue dots). The IPD ratio cutoff was marked by dashed line. **B.** Correlation of penetrance in gradient-partitioned CCS datasets of *Escherichia coli* at varying coverages. The full 42× dataset was subdivided into 21 partial datasets, each with 2× decremental coverage. Spearman correlation coefficient was calculated between each pair of datasets. **C.** Correlation of ΣP (the sum of penetrance within 50, 100, 200, 500 and 1,000 bp of the reference genome) in gradient-partitioned CCS datasets of *E. coli* with different coverages. The full 42× dataset was partitioned into 8 partial datasets with 5× descending per dataset. Spearman correlation coefficient was calculated between each pair of datasets.

For organisms with low 6mA levels, such as most multicellular eukaryotes (e.g., *Homo sapiens*, *Arabidopsis thaliana*, *Drosophila melanogaster*, and *Caenorhabditis elegans*), SMAC is unable to identify 6mA due to the absence of a bimodal distribution in their IPD ratios (Figure S3). In contrast, SMAC can successfully detect 6mA in prokaryotes and lower unicellular eukaryotes, such as *E. coli*, *Chlamydomonas reinhardtii* (Figure S3) and *T. thermophila*. Notably, for organisms with high 6mApT abundance, SMAC offers a more precise determination of 6mApT states (Figure 4, Table S([0-9]+)).

At the single-molecule level, SMAC can provide reliable 6mA detection without requiring high sequencing coverage. However, at the genome-wide level, a certain sequencing depth is necessary. We performed a gradient partition of an *E. coli* CCS dataset with approximately 42× depth and used SMAC to calculate the penetrance of individual A sites and the 6mA level (ΣP, the sum of penetrance within a specific region of the reference genome). The correlation of penetrance between partial dataset and full dataset gradually increased and stabilized as the coverage of the partial dataset approached that of the full dataset, with the Spearman correlation coefficient surpassing 0.9 once the depth exceeded 20× (Figure 5B). A similar pattern is observed with ΣP (Figure 5C). Additionally, higher sequencing depth could reduce the discrepancy on penetrance and methylation level between SMAC and standard SMRT-seq pipelines (Figure S4). Therefore, SMAC recommends a minimal sequencing depth of over 20× coverage for genome-wide 6mA assessment.

### Conclusion and perspective

In this work, we developed SMAC, an automated framework for detecting 6mA based on SMRT CCS data. SMAC applies rigorous data pre-processing with adjustable parameters, allowing users to balance data yield and quality according to their specific needs. By analyzing molecule-specific IPD ratios and leveraging the statistical distribution characteristics of these ratios, SMAC identifies 6mA at single-molecule, single-nucleotide resolution with high confidence. It detects low penetrance sites and uncovers hidden 6mA patterns at 6mApT dinucleotides that may be overlooked or misinterpreted in studies using the standard SMRT-seq pipeline.

The demarcation of 6mA states by SMAC provides the resolution needed to study the catalytic features of 6mA methyltransferase (MTase) (38). SMAC is essential in revealing that the catalytic product of the *de novo* MTase AMT2 is hemi-6mApT (39). Additionally, by tracing cell cycle-dependent 6mA dynamics, SMAC uncovers the hemi-to-full conversion mediated by the maintenance MTase AMT1 (10), a discovery impossible with CLR-based ensemble analysis (7).

SMAC can also empower us to explore 6mA functions during environmental responses. In the ciliate *Pseudocohnilembus persalinus* and the fungus *Phycomyces blakesleeanus*, symmetric and asymmetric methylated 6mApT have been demonstrated to fulfill different biological roles (12,32). However, both studies relied on the CLR data analyzed by the standard SMRT-seq pipeline (12,32). It is conceivable that SMAC could offer more precise determination of hemi- and full-methylation in *P. persalinus*, *P. blakesleeanus*, and other species, thereby enhancing our understanding of the biological significance of differentially methylated 6mA.

The PacBio Sequel II system offers significantly higher data output compared to its predecessor, the RS II system, making it the platform of choice in most studies utilizing PacBio sequencing. SMAC is specifically designed to process subreads data generated from the Sequel II system. However, with the advancements in PacBio’s sequencing technology, the recently introduced Revio system delivers even greater data yields than Sequel II. As a result, future adjustment and optimization to our workflow may be necessary to accommodate the increased data capacity of the Revio system.

## Data availability

CCS datasets of were downloaded from the NCBI database with the following BioProject accession numbers: *Tetrahymena thermophila* (PRJNA932808), *Caenorhabditis elegans* (PRJNA857919) and various other organisms including human PBMC (peripheral blood mononuclear cells), *Arabidopsis thaliana*, *Drosophila melanogaster*, *Escherichia coli*, *Chlamydomonas reinhardtii* and a mixed sample containing *Escherichia coli*, *Helicobacter pylori* and *Saccharomyces cerevisiae* (PRJNA667898). The SMAC pipeline is available along with a tutorial at GitHub (https://github.com/liiihc/SMAC).

## Author contribution

Haicheng Li, Junhua Niu and Yalan Sheng: methodology, formal analysis, investigation, writing - original draft, writing - review & editing. Yifan Liu and Shan Gao: conceptualization, supervision, funding acquisition, writing-review & editing.

## Acknowledgements

High-performance computing resources for data processing were provided by the Institute of Evolution and Marine Biodiversity at OUC, High-Performance Biological Supercomputing Center at the Ocean University of China for this research, Marine Big Data Center of Institute for Advanced Ocean Study at OUC, and the Center for Advanced Research Computing (CARC) at the University of Southern California. Our special thanks are given to Weibo Song (OUC) for his helpful suggestions during drafting the manuscript.

## Figure legend

**Fig. S1.**
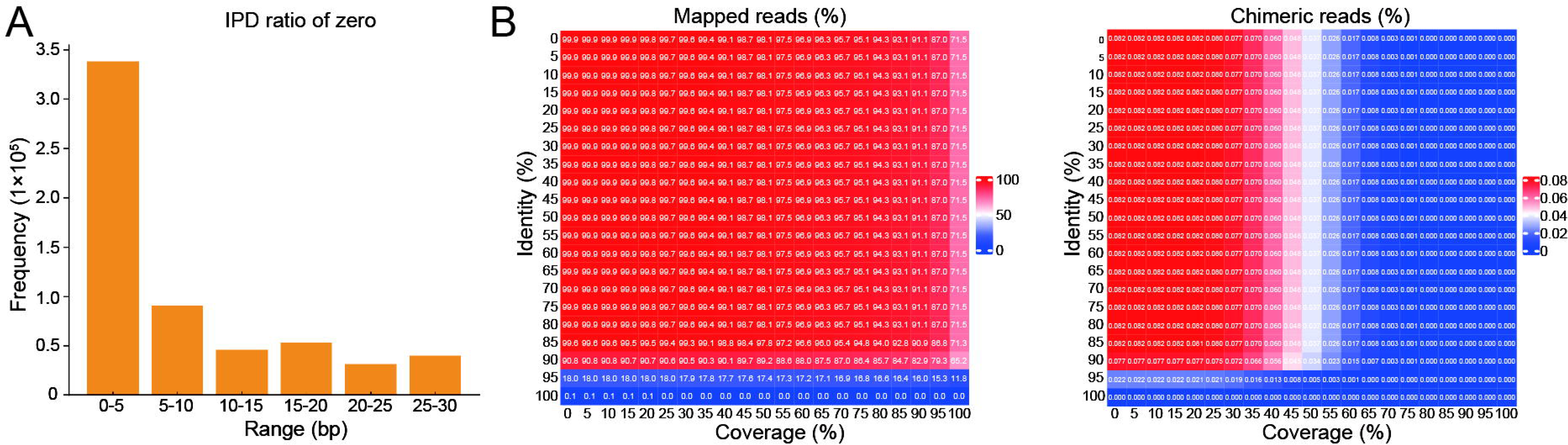
Base trimming near adapters and filtration of contamination reads by BLASTN. **A.** Distribution of abnormal bases with an IPD ratio of zero within 0-5, 5-10, 10-15, 15-20, 20-25, and 25-30 bp near adapters. **B.** Effect of coverage (X-axis) and identity (Y-axis) in the BLASTN alignment on reads ratio (left) and chimeric reads ratio (right), shown as a heatmap.

**Fig. S2.**
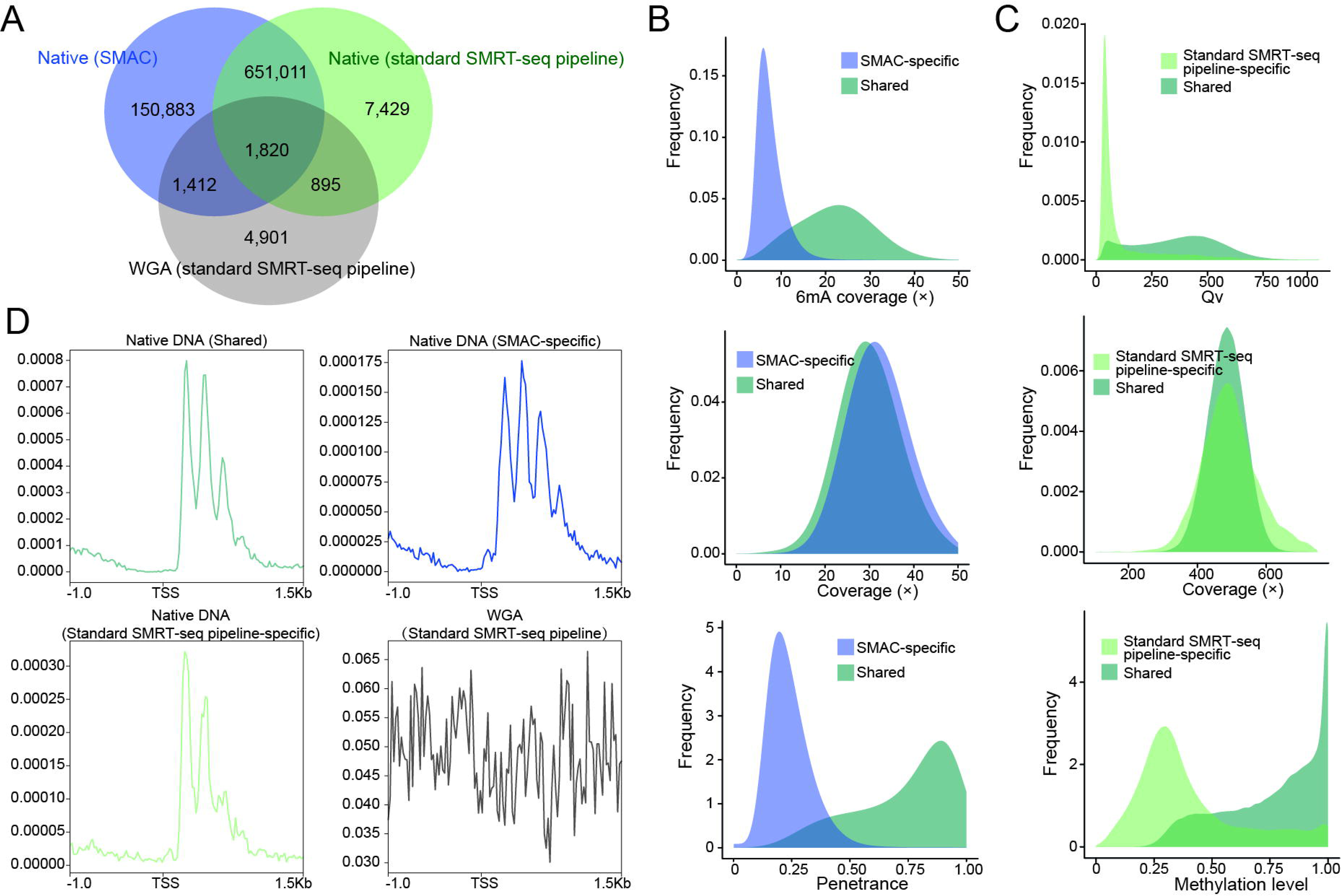
Comparation of SMAC and standard SMRT-seq pipeline results. **A.** Overlap of 6mA sites identified by SMAC in native DNA sample (6mA coverage ≥ 5) with sites identified by the standard SMRT-seq pipeline in native DNA (coverage ≥ 100, Qv ≥ 30) and in WGA (coverage ≥ 100, Qv ≥ 30). **B.** Distribution of 6mA coverage (sequencing depth), coverage (sequencing depth) and penetrance calculated by SMAC. 6mA sites were divided into SMAC-specific group and shared group between SMAC and standard SMRT-seq pipeline. **C.** Qv, coverage (sequencing depth * passes) and methylation level calculated by standard SMRT-seq pipeline in native DNA. 6mA sites were divided into standard SMRT-seq pipeline-specific group and shared group between SMAC and standard SMRT-seq pipeline. **D.** Distribution profiles of 10,000 randomly selected SMAC-specific 6mA sites in native DNA sample (blue line), 8,324 standard SMRT-seq pipeline-specific 6mA sites in native DNA sample (light green line), 10,000 randomly selected shared 6mA sites in native DNA sample (dark green line), as well as 9,024 standard SMRT-seq pipeline defined false positive sites in WGA (gray line). 6mA peaks had a periodicity distribution downstream of TSS, except for those false-positive sites in WGA.

**Fig. S3.**
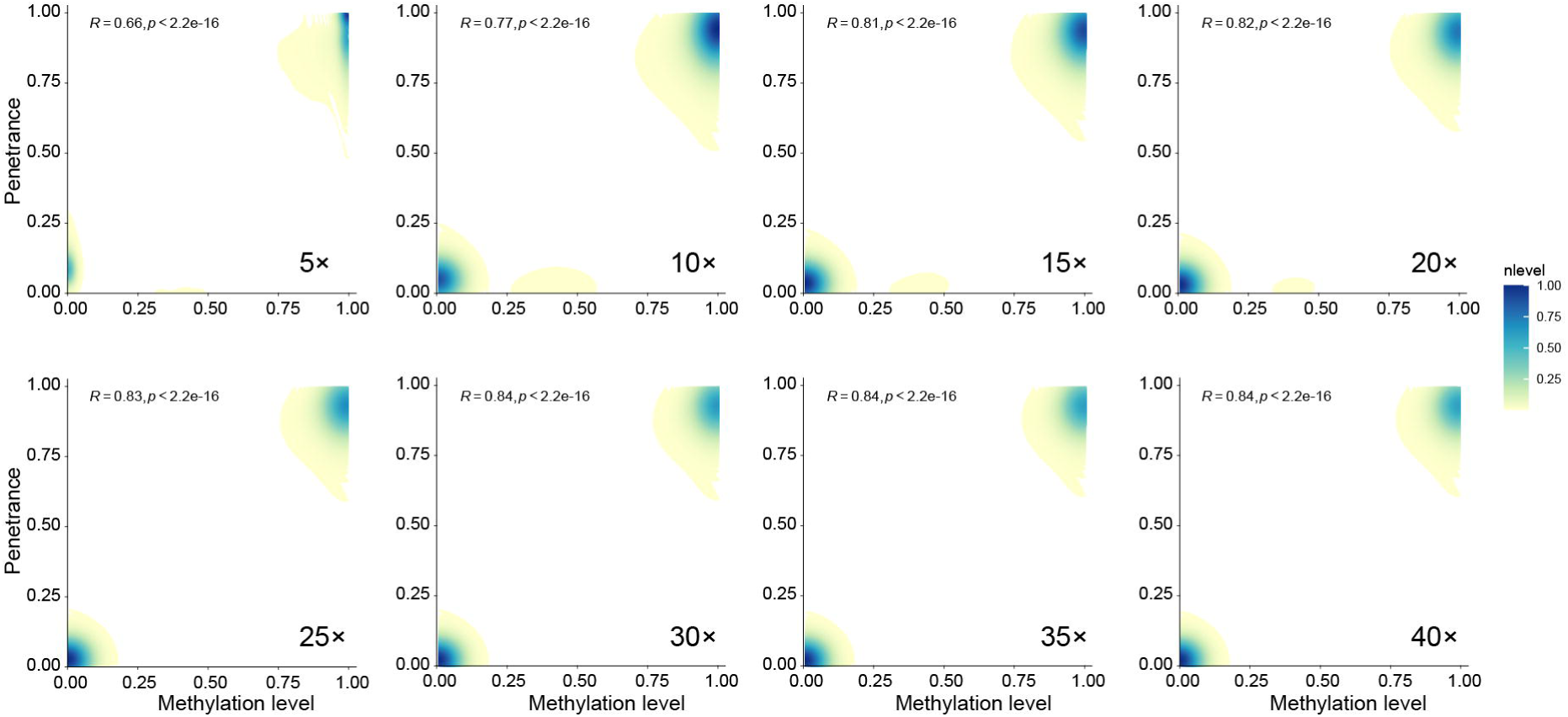
Performance of SMAC in high-6mA and low-6mA samples. IPD ratio distribution (Log_2_) of A sites in low-6mA samples (*Homo sapiens*, *Arabidopsis thaliana*, *Drosophila melanogaster*, and *Caenorhabditis elegans*), and deconvolution of the 6mA peak and the unmodified A peak for IPD ratio distribution in high-6mA samples (*E. coli* and *Chlamydomonas reinhardtii*). Residual (light blue dots) represented the difference between authentic IPD distribution and the best fit curve (red dots). The IPD ratio cutoff was marked by dashed line.

**Fig. S4.**
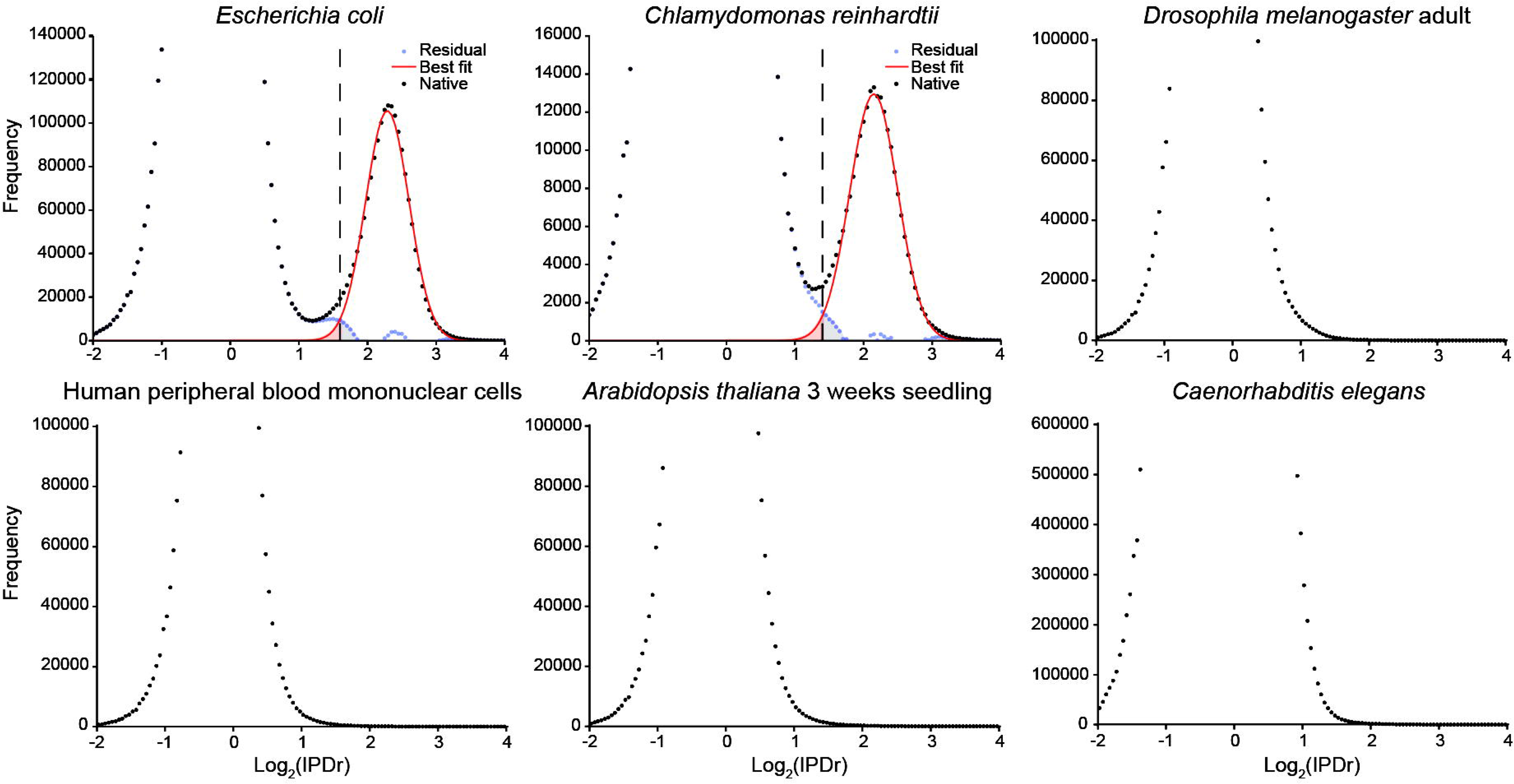
Relationship between sequencing coverage, penetrance, and methylation level. Correlation of penetrance calculated by SMAC and methylation level calculated by the standard SMRT-seq pipeline in gradient-partitioned CCS datasets of *Escherichia coli* with varying coverages. The full 42× dataset was partitioned into datasets with coverages of 40×, 35×, 30×, 25×, 20×, 15×, 10×, and 5×. 6mA site distribution density was plotted as a heat map.

**Table S1.** Motif distribution for 6mA dinucleotide.

| **Motif** | **Native DNA**<br>**shared** | **Native DNA**<br>**SMAC-specific** | **Native DNA**<br>**ensemble-specific** | **WGA ensemble** |
| --- | --- | --- | --- | --- |
| AT | 99.55% | 92.20% | 75.65% | 13.65% |
| AA | 0.10% | 2.85% | 6.92% | 22.51% |
| AC | 0.13% | 1.69% | 8.30% | 28.27% |
| AG | 0.22% | 3.26% | 9.13% | 35.57% |
| total | 100.00% | 100.00% | 100.00% | 100.00% |

**Table S2.** *Tetrahymena* strains used in this study (1,2).

| Strains | 6mA<sub>P</sub>T states | Site number | Percentage (%) |
| --- | --- | --- | --- |
| Wild type | Symmetric | 616,496 | 93.95 |
|  | Asymmetric-W | 19,871 | 3.03 |
|  | Asymmetric-C | 19,801 | 3.02 |
|  | Full | 15,300,728 | 89.29 |
|  | Hemi-W | 915,693 | 5.34 |
|  | Hemi-C | 919,715 | 5.37 |
| ΔAMT1 | Symmetric | 17,904 | 12.29 |
|  | Asymmetric-W | 64,002 | 43.93 |
|  | Asymmetric-C | 63,789 | 43.78 |
|  | Full | 73,346 | 2.37 |
|  | Hemi-W | 1,512,272 | 48.83 |
|  | Hemi-C | 1,511,401 | 48.80 |

## References

1. Fu, Y., Luo, G.-Z., Chen, K., Deng, X., Yu, M., Han, D., Hao, Z., Liu, J., Lu, X., Doré, Louis C. et al. (2015) N^6^-methyldeoxyadenosine marks active transcription start sites in *Chlamydomonas*. Cell, 161, 879–892.

2. Zhang, G.Q., Huang, H., Liu, D., Cheng, Y., Liu, X.L., Zhang, W.X., Yin, R.C., Zhang, D.P., Zhang, P., Liu, J.Z. et al. (2015) N^6^-methyladenine DNA modification in *Drosophila*. Cell, 161, 893–906.

3. Wu, T.P., Wang, T., Seetin, M.G., Lai, Y., Zhu, S., Lin, K., Liu, Y., Byrum, S.D., Mackintosh, S.G., Zhong, M. et al. (2016) DNA methylation on N^6^-adenine in mammalian embryonic stem cells. Nature, 532, 329–333.

4. Wang, Y., Chen, X., Sheng, Y., Liu, Y. and Gao, S. (2017) N^6^-adenine DNA methylation is associated with the linker DNA of H2A.Z-containing well-positioned nucleosomes in Pol II-transcribed genes in *Tetrahymena*. Nucleic Acids Research, 45, 11594–11606.

5. Xie, Q., Wu, T.P., Gimple, R.C., Li, Z., Prager, B.C., Wu, Q., Yu, Y., Wang, P., Wang, Y., Gorkin, D.U. et al. (2018) N^6^-methyladenine DNA Modification in Glioblastoma. Cell, 175, 1228–1243.e1220.

6. Beh, L.Y., Debelouchina, G.T., Clay, D.M., Thompson, R.E., Lindblad, K.A., Hutton, E.R., Bracht, J.R., Sebra, R.P., Muir, T.W. and Landweber, L.F. (2019) Identification of a DNA N^6^-Adenine Methyltransferase Complex and Its Impact on Chromatin Organization. Cell, 177, 1781–1796.e1725.

7. Wang, Y., Sheng, Y., Liu, Y., Zhang, W., Cheng, T., Duan, L., Pan, B., Qiao, Y., Liu, Y. and Gao, S. (2019) A distinct class of eukaryotic MT-A70 methyltransferases maintain symmetric DNA N^6^-adenine methylation at the ApT dinucleotides as an epigenetic mark associated with transcription. Nucleic Acids Research, 47, 11771–11789.

8. Li, Z., Zhao, S., Nelakanti, R.V., Lin, K., Wu, T.P., Alderman, M.H., Guo, C., Wang, P., Zhang, M., Min, W. et al. (2020) N^6^-methyladenine in DNA antagonizes SATB1 in early development. Nature, 583, 625–630.

9. Zhang, S., Li, B., Du, K., Liang, T., Dai, M., Huang, W., Zhang, H., Ling, Y. and Zhang, H. (2020) Epigenetically modified N^6^-methyladenine inhibits DNA replication by human DNA polymerase iota. Biochimie, 168, 134–143.

10. Sheng, Y., Wang, Y., Yang, W., Wang, X.Q., Lu, J., Pan, B., Nan, B., Liu, Y., Ye, F., Li, C. et al. (2024) Semiconservative transmission of DNA N^6^-adenine methylation in a unicellular eukaryote. Genome Research, 34, 740–756.

11. Sheng, Y., Pan, B., Wei, F., Wang, Y., Gao, S. and Yao, M.-C. (2021) Case study of the response of N^6^-methyladenine DNA modification to environmental stressors in the unicellular eukaryote *Tetrahymena thermophila*. mSphere, 6, e01208–01220.

12. Liu, Y., Niu, J., Ye, F., Solberg, T., Lu, B., Wang, C., Nowacki, M. and Gao, S. (2024) Dynamic DNA N^6^-adenine methylation (6mA) governs the encystment process, showcased in the unicellular eukaryote *Pseudocohnilembus persalinu*. Genome Research, 34, 256–271.

13. Boulias, K. and Greer, E.L. (2022) Means, mechanisms and consequences of adenine methylation in DNA. Nature Reviews Genetics, 23(7), 411–428.

14. Feng, X. and He, C. (2023) Mammalian DNA N^6^-methyladenosine: Challenges and new insights. Molecular Cell, 83, 343–351.

15. Lyu, C., Wang, H.-D., Lai, W. and Wang, H. (2023) Identification and quantification of DNA N^6^-methyladenine modification in mammals: A challenge to modern analytical technologies. Current Opinion in Chemical Biology, 73, 102259.

16. Eid, J., Fehr, A., Gray, J., Luong, K., Lyle, J., Otto, G., Peluso, P., Rank, D., Baybayan, P., Bettman, B. et al. (2009) Real-Time DNA Sequencing from Single Polymerase Molecules. Science, 323, 133–138.

17. Flusberg, B.A., Webster, D.R., Lee, J.H., Travers, K.J., Olivares, E.C., Clark, T.A., Korlach, J. and Turner, S.W. (2010) Direct detection of DNA methylation during single-molecule, real-time sequencing. Nature Methods, 7, 461–465.

18. Fang, G., Munera, D., Friedman, D.I., Mandlik, A., Chao, M.C., Banerjee, O., Feng, Z., Losic, B., Mahajan, M.C., Jabado, O.J. et al. (2012) Genome-wide mapping of methylated adenine residues in pathogenic *Escherichia coli* using single-molecule real-time sequencing. Nature Biotechnology, 30, 1232–1239.

19. Roberts, R.J., Carneiro, M.O. and Schatz, M.C. (2013) The advantages of SMRT sequencing. Genome Biology, 14, 405.

20. Schadt, E.E., Banerjee, O., Fang, G., Feng, Z., Wong, W.H., Zhang, X., Kislyuk, A., Clark, T.A., Luong, K., Keren-Paz, A. et al. (2013) Modeling kinetic rate variation in third generation DNA sequencing data to detect putative modifications to DNA bases. Genome Research, 23, 129–141.

21. Rhoads, A. and Au, K.F. (2015) PacBio Sequencing and Its Applications. Genomics, Proteomics & Bioinformatics, 13, 278–289.

22. Ardui, S., Ameur, A., Vermeesch, J.R. and Hestand, M.S. (2018) Single molecule real-time (SMRT) sequencing comes of age: applications and utilities for medical diagnostics. Nucleic Acids Research, 46, 2159–2168.

23. Beaulaurier, J., Zhang, X.-S., Zhu, S., Sebra, R., Rosenbluh, C., Deikus, G., Shen, N., Munera, D., Waldor, M.K., Chess, A. et al. (2015) Single molecule-level detection and long read-based phasing of epigenetic variations in bacterial methylomes. Nature Communications, 6, 7438.

24. Abdulhay, N.J., McNally, C.P., Hsieh, L.J., Kasinathan, S., Keith, A., Estes, L.S., Karimzadeh, M., Underwood, J.G., Goodarzi, H., Narlikar, G.J. et al. (2020) Massively multiplex single-molecule oligonucleosome footprinting. eLife, 9, e59404.

25. Stergachis, A.B., Debo, B.M., Haugen, E., Churchman, L.S. and Stamatoyannopoulos, J.A. (2020) Single-molecule regulatory architectures captured by chromatin fiber sequencing. Science, 368, 1449–1454.

26. Feng, Z., Fang, G., Korlach, J., Clark, T., Luong, K., Zhang, X., Wong, W. and Schadt, E. (2013) Detecting DNA Modifications from SMRT Sequencing Data by Modeling Sequence Context Dependence of Polymerase Kinetic. PLOS Computational Biology, 9, e1002935.

27. Kong, Y., Cao, L., Deikus, G., Fan, Y., Mead, E.A., Lai, W., Zhang, Y., Yong, R., Sebra, R., Wang, H. et al. (2022) Critical assessment of DNA adenine methylation in eukaryotes using quantitative deconvolution. Science, 375, 515–522.

28. Zhang, J., Peng, Q., Ma, C., Wang, J., Xiao, C., Li, T., Liu, X., Zhou, L., Xu, X., Zhou, W.-Z. et al. (2023) 6mA-Sniper: Quantifying 6mA sites in eukaryotes at single-nucleotide resolution. Science Advances, 9, eadh7912.

29. Liang, Z., Shen, L., Cui, X., Bao, S., Geng, Y., Yu, G., Liang, F., Xie, S., Lu, T., Gu, X. et al. (2018) DNA N^6^-Adenine Methylation in *Arabidopsis thaliana*. Developmental Cell, 45, 406–416.e403.

30. Zhang, Q., Liang, Z., Cui, X., Ji, C., Li, Y., Zhang, P., Liu, J., Riaz, A., Yao, P., Liu, M. et al. (2018) N6-Methyladenine DNA Methylation in *Japonica* and *Indica* Rice Genomes and Its Association with Gene Expression, Plant Development, and Stress Responses. Molecular Plant, 11, 1492–1508.

31. Zhou, C., Wang, C., Liu, H., Zhou, Q., Liu, Q., Guo, Y., Peng, T., Song, J., Zhang, J., Chen, L. et al. (2018) Identification and analysis of adenine N^6^-methylation sites in the rice genome. Nature Plants, 4, 554–563.

32. Lax, C., Mondo, S.J., Osorio-Concepción, M., Muszewska, A., Corrochano-Luque, M., Gutiérrez, G., Riley, R., Lipzen, A., Guo, J., Hundley, H. et al. (2024) Symmetric and asymmetric DNA N^6^-adenine methylation regulates different biological responses in Mucorales. Nature Communications, 15, 6066.

33. Hahn, A., Hung, G.C.C., Ahier, A., Dai, C.-Y., Kirmes, I., Forde, B.M., Campbell, D., Lee, R.S.Y., Sucic, J., Onraet, T. et al. (2024) Misregulation of mitochondrial 6mA promotes the propagation of mutant mtDNA and causes aging in *C. elegans*. Cell Metabolism.

34. Zhao, L., Gao, F., Gao, S., Liang, Y., Long, H., Lv, Z., Su, Y., Ye, N., Zhang, L., Zhao, C. et al. (2021) Biodiversity-based development and evolution: the emerging research systems in model and non-model organisms. SCIENCE CHINA Life Sciences, 64, 1236–1280.

35. Ye, F., Chen, X., Ju, A., Sheng, Y., Duan, L., Al-Rasheid, K.A.S., Stover, N.A. and Gao, S. (2024) Comprehensive genome annotation of the model ciliate *Tetrahymena thermophila* by in-depth epigenetic and transcriptomic profiling. bioRxiv, 2024.2001.2031.578305.

36. Pan, B., Ye, F., Li, T., Wei, F., Warren, A., Wang, Y. and Gao, S. (2023) Potential role of N^6^-adenine DNA methylation in alternative splicing and endosymbiosis in *Paramecium bursaria*. iScience, 26, 106676.

37. Zhao, H., Ma, J., Tang, Y., Ma, X., Li, J., Li, H. and Liu, Z. (2024) Genome-wide DNA N^6^-methyladenosine in *Aeromonas veronii* and *Helicobacter pylori*. BMC Genomics, 25, 161.

38. Wang, Y., Nan, B., Ye, F., Zhang, Z., Yang, W., Pan, B., Niu, J., Ju, A., Liu, Y., Zhang, W. et al. (2024) Dual modes of DNA N^6^-methyladenine maintenance by distinct methyltransferase complexes. bioRxiv, 2024.2007.2021.604504.

39. Cheng, T., Zhang, J., Li, H., Diao, J., Zhang, W., Niu, J., Kataoka, K. and Gao, S. (2024) Identification and characterization of the *de novo* methyltransferases for eukaryotic N^6^-methyladenine (6mA). bioRxiv, 2024.2003.2025.586193.

